# Technical advances in the development of zonation liver *in vitro* systems that incorporate localized Wnt activating signals

**DOI:** 10.1101/2021.04.01.438073

**Authors:** Eider Valle-Encinas, Michael Dawes, Carmen Velasco Martinez, Kate McSweeney, Miryam Müller, Tom Bird, Trevor Dale

## Abstract

A Wnt microenvironment sustained by the hepatic central vein is essential for the segregation of liver functions into zones. Current liver culture systems lack localized Wnt cues and as a consequence fail to maintain the hepatocyte functional heterogeneity that is observed in the intact organ. In this study, organoid models and 2D-culture systems were used to identify cellular sources and Wnt presentation methods that could support the future development of zonated liver *in vitro* systems. Using soluble ligands, we show that primary hepatocyte (PH)-derived organoids but not bile duct (BD)-derived organoids may be used to recapitulate the resting liver. We provide evidence that differentiation of PH-organoids in the presence of Wnt9b and Rspo3 induce pericentral maturation. Finally, we show that immobilization of Rspo3 onto beads in combination with soluble Wnt9b may be a valid strategy to recreate the central vein Wnt microenvironment *in vitro*.

## Introduction

With an estimated development cost of $2.6 billion per drug that reaches the clinic, there is a need for *in vitro* liver systems that can reduce late stage failure of candidate drugs due to hepatotoxicity ^1^. Accurate modelling the in vivo liver responses *remains* challenging because hepatocytes display tremendous functional heterogeneity that is spatially segregated in the smallest functional unit of the liver; the hepatic lobule ^2, 3^. Based on the position in this unit, the hepatocytes can be broadly clustered in two main functional pools: pericentral and periportal ^2–4^. Periportal hepatocytes are located in the periphery of each lobular unit; they receive nutrient- and oxygen-rich blood from the digestive tract through the portal triad and are overall more invested in the synthesis of albumin, gluconeogenesis and lipid *β*-oxidation ^3, 5–8^ ^9^. By contrast, pericentral hepatocytes, located in the center, receive blood-transported metabolites secreted by upstream periportal hepatocytes and are more invested in the production of bile, glycolysis and the synthesis of glutamine ^3, 5, 6, 8, 10^. The expression of drug metabolism enzymes as well as efflux basolateral transporters is also zonated, with enzymes from the cytochrome P450 superfamily especially enriched in pericentral hepatocytes ^8, 11^. The fine-coupling of metabolic processes between neighboring hepatocytes presents a major challenge for those attempting to accurately recapitulate liver responses *in vitro*, since culture systems need to recapitulate hepatocyte functional heterogeneity and hepatocyte-to-hepatocyte communication and metabolic zonation in a continuous.

Following injury, hepatocytes derived from a one lobular region are able to repair damage and to subsequently acquire appropriate zonal characteristics, implying that metabolic zonation is a primarily driven by cell extrinsic ‘niche’ factors ^12–14^. Jungermann and Katz proposed that graded oxygen, nutrients, hormone and metabolite gradients from the polarized bloodstream of the lobule were key determinants of zonation ^4^. An important role for the canonical Wnt pathway in pericentral hepatocytes has been shown for the expression of pericentral genes and conversely, for the repression of periportal markers in pericentral lobular areas ^2, 15–20^. The canonical Wnt pathway is initiated through the binding of Wnt proteins to a receptor from the Fzd family (Fzd1-10) and a receptor from the Lrp family (Lrp5 or Lrp6); leading to the activation of a signaling pathway that culminates in the stabilization and nuclear translocation of the transcriptional coactivator *β*-catenin ^2^. Asymmetric activation of the Wnt/*β*-catenin pathway in the hepatic lobule has been attributed to the secretion of Wnt proteins (Wnt2 and Wnt9b) by endothelial cells of the hepatic central vein, although direct evidence of the importance of these two ligands in the modulation of zonation is missing ^2, 15, 20–22^. Central vein endothelial cells additionally produced Rspo3, a Wnt agonist that potentiates the activity of Wnt proteins by stabilizing Wnt receptors and that was shown to be critical for the establishment of a pericentral metabolic program ^19, 23^.

Synthetic reconstruction of zonation *in vitro* will require the incorporation of cell extrinsic factors that dictate zonal patterns. While previous work investigated the incorporation of oxygen, metabolite and hormone gradients within liver *in vitro* systems, to our knowledge, only one study by Whalicht et al. (2020) has attempted to recapitulate Wnt-driven zonal heterogeneity to date ^24–31^. Whalicht et al (2020) used immortalized murine hepatocyte cultures transduced with a Tet-On-*Δβ*cat synthetic construct in which the expression of a mutant active form of *β*-catenin was induced following doxycycline administration to induce the periportal- and pericentral-like hepatocyte gene expression signatures ^31^. However, there were significant disadvantages associated with this strategy including the use of transformed hepatocytes, the fact that b-catenin activation is unlikely to be completely equivalent to Wnt pathway activation by endogenous ligands and that *β*-catenin overexpression was randomization between cells, precluding the spatial organization of zonation at scales of 10-20 cell diameters that is observed in the lobular unit *in vivo*.

In addition to the integration of zonal microenvironments, a critical component for the reconstruction of zonation *in vitro* will be a renewable source of cells that can accurately recapitulate the responses of mature hepatocytes. Two types of hepatic organoids derived from adult tissue have recently been described that allow long-term culture ^32–34^. Bile duct-derived organoids (BD organoids) are 3D structures derived from biliary epithelial cells (BECs; also known as cholangiocytes); that can be differentiated into hepatocyte-like cells when exposed to a defined cocktail of growth factors including the Wnt pathway modulator Rspo1 ^32^. By contrast, primary hepatocyte-derived organoids (PH organoids) are maintained in a distinct cocktail of growth factors including TNF*α* and CHIR99021, a GSK3 inhibitor that activates the Wnt/*β*-catenin pathway ^34 33^. We recently showed that, despite of the growing popularity of BD organoids as cellular platform for hepatotoxicity assays, BD organoid cultures do not capture the biology of mature resting hepatocytes and that, by contrast with Whalicht et al. (2020) in immortalized hepatocytes, synthetic stabilization of *β*-catenin was unable to induce the expression of pericentral metabolic genes ^31, 35^.

In the present study, we report significant advances in the development of liver *in vitro* systems with features of the zonated liver. First, we evaluated the competency of PH and BD organoids as cellular source to model zonation by assessing their response to purified central vein Wnt activators that are now available (Wnt9b and Rspo3 proteins). Our data strongly supports the use of PH but not BD organoids as a cellular source for the synthetic reconstruction of liver zonation. We show that differentiation of PH organoids in the presence of Wnt9b and Rspo3 induced a more specific pericentral transcriptional program than the currently available CHIR-based protocols. Using HEK293T Wnt-reporter cell lines we further demonstrate that the covalent immobilization of Rspo3 in combination with soluble Wnt9b may be a valid strategy to recreate a central vein microenvironment *in vitro*.. When combined, these findings open new frontiers in the development of liver *in vitro* systems with features of the zonated liver.

## Results and Discussion

### PH organoids but not BD organoids mirror the response of primary hepatocyte 2D cultures to Wnt9b and Rspo3

The Wnt/*β*-catenin pathway regulates the expression of *Axin2*, a well-established *β*-catenin target across tissues, and the metabolic genes *Cyp1a2*, *Cyp2e1* and *Glul* (gene coding for GS) in resting pericentral hepatocytes. To assess the competency of PH and BD organoids as a cell sources to reproduce Wnt-driven zonation *in vitro*, the expression of *Axin2, Cyp1a2, Cyp2e1* and *Glul* was assessed by RT-qPCR following exposure to Wnt9b and Rspo3, two Wnt pathway upstream activators secreted by the central vein. Prior to Wnt9b and Rspo3 treatment, both BD and PH organoids were taken through their respective ‘differentiation’ protocols as previously described and as summarized in the timelines shown (Figure 1) ^32, 33^. The responses of PH and BD organoids were compared to those of freshly-isolated primary hepatocytes cultured as conventional monolayers on collagen-coated plates (Figure 1, A).

**Figure 1.**
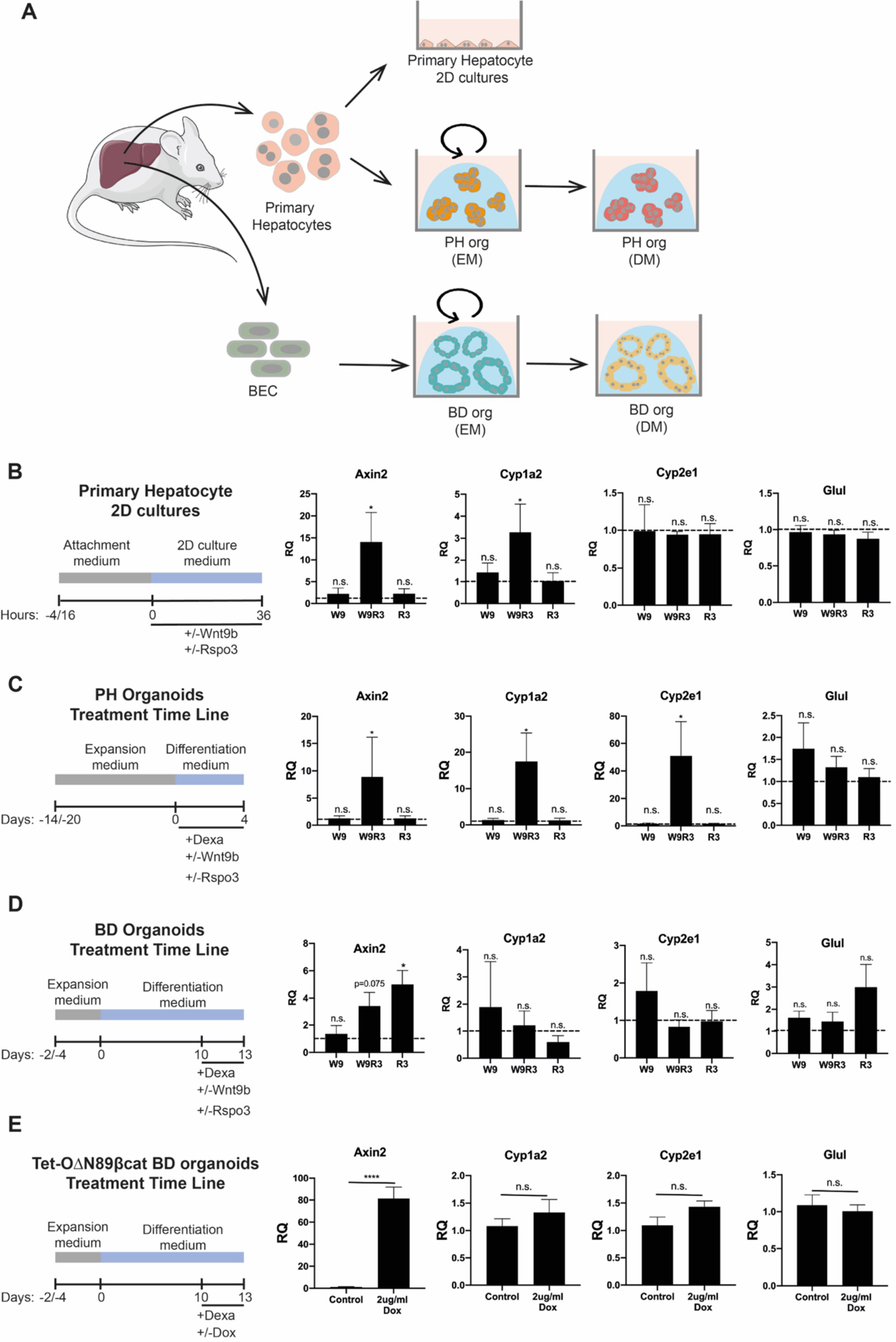
(A) Scheme describing the process of establishment of the primary hepatocyte, BD organoids and PH organoids cultures used in this study. (B-D) RT-qPCR gene expression analysis of *β*-catenin target genes in the liver in wild-type primary hepatocyte 2D cultures (panel A), PH organoids (panel B) and BD organoids (panel C) exposed to Wnt9b (100ng/ml) or Rspo3 (50ng/ml). Treatment time line for each culture type is specified on the left panel. For hepatocyte (n=4 isolations from different animals). For PH organoid samples, n=4 (Table S3). For BD organoids samples, n=5 (Table S3). (D) RT-qPCR gene expression analysis in Tet-O*Δ*N89*β*cat BD organoids upon activation of mutant *β*-catenin following exposure to doxycycline (Dox) at a concentration of 2 mg/ml. Treatment time line is shown on the left panel. n=9 (Table S3).

Mirroring the responses of the primary hepatocyte 2D cultures, concomitant exposure to Wnt9b and Rspo3, but not the individual ligands, significantly increased Axin2 and Cyp1a2 levels in PH organoids (Compare Figure 1 B and C). Cyp1a2 was more strongly induced in PH organoids than in the primary hepatocyte monolayers and Cyp2e1 was only induced in PH organoids (Figure 1 B and C). GluI was not induced by Wnt9b and Rspo3 in either primary hepatocytes or PH organoids, suggesting that additional features of *in vivo* biology remain to be recapitulated *in vitro* (Figure 1 B and C). These data were consistent with the findings of Wahlicht et al. (2020) similarly saw no induction of Glul expression following synthetic stabilization of β-catenin in immortalized mouse hepatocytes ^31^.

The classic Wnt target gene Axin2 was induced by Rspo3 alone and in combination with Wnt9b in BD organoids, indicating that exposure to Rspo3 was sufficient to activate the Wnt/*β*-catenin in BD organoids (Figure1 D). The increase in Axin2 mRNA levels in BD organoids was, however, not associated with an increase in expression of the pericentral metabolic genes Cyp1a2 and Cyp2e1 (Figure1 D) ^35^. To exclude the possibility that BD organoids may fail to express the requisite receptors to respond to the Wnt ligands, we derived BD organoids from a Tet-O-ΔN89β-catenin mouse model that supports the doxycycline-induced stabilization of *β*-catenin ^35, 36^. Following induction of *β*-catenin with 2 µg/ml of doxycycline during the last three days of BD organoid differentiation, Axin2 was strongly induced, but none of the pericentral metabolic genes responded (Figure1, E). These results are consistent with our previous findings using lower concentrations of doxycycline (0.1 µg/ml) and show that while BD organoids are responsive to Wnt activation and having undergone some aspects of hepatocyte differentiation, cannot be used to recapitulate zonation *in vitro*

In the adult liver, the Wnt/*β*-catenin pathway controls hepatocyte proliferation and regeneration following injury. We previously showed that stabilization of β-catenin with 0.1µg/ml of doxycycline in differentiated Tet-O-ΔN89β-catenin BD organoids caused an increase in Hepatocyte Progenitor Cell (HPC) gene expression ^35^. To determine whether Wnt9b and Rspo3 elicited a similar transcriptional program in PH organoids, the expression of Lgr5, Sox9, Spp1, CyclinD1 and Mki67 was examined in differentiated cultures. With the exception of Lgr5 none of the genes evaluated (Sox9, Spp1, Cyclin D1 and Mki67) were significantly induced in primary hepatocyte cultures or in PH organoid cultures (Figure 2 A and B). This response contrasted with the HPC program of Tet-O-ΔN89β-catenin BD organoids suggesting that the organoids recapitulate distinct cellular contexts (Figure 2 C).

**Figure 2.**
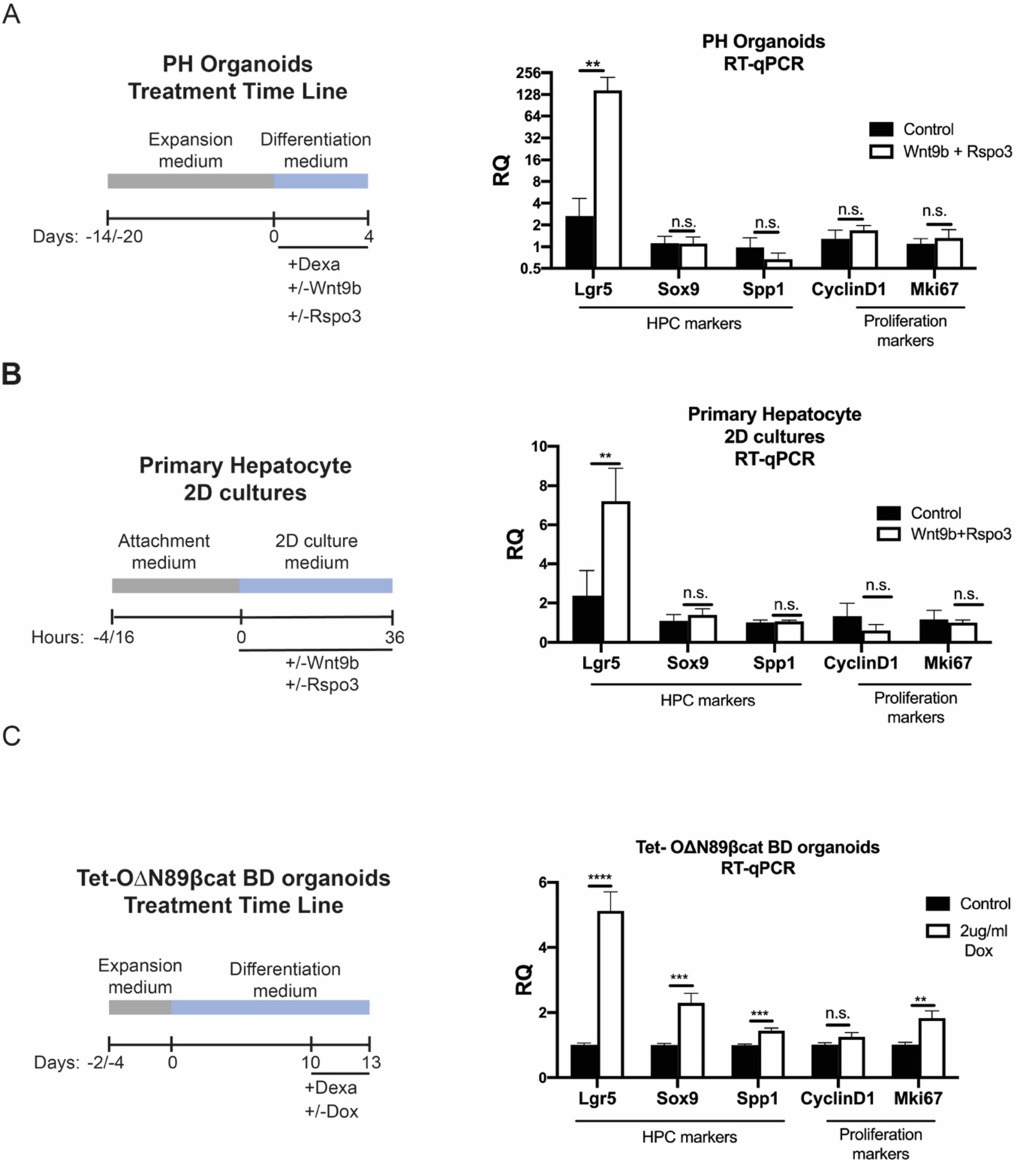
(A) RT-qPCR gene expression analysis in wild-type PH 2D cultures following activation of the Wnt/*β*-catenin pathway with Wnt9b (100ng/ml) and Rspo3 (50ng/ml) (n=3 isolations from different animals). (B) RT-qPCR gene expression analysis of PH organoids where the Wnt/*β*-catenin pathway was activated following exposure with Wnt9b (100ng/ml) and Rspo3 (50ng/ml). Left panel indicates treatment timeline. n=4 (Table S3). (C) RT-qPCR gene expression analysis in Tet-O*Δ*N89*β*cat BD organoids upon activation of the Wnt/*β*-catenin pathway with 2 µg/ml of doxycycline (Dox). Treatment time line is shown in the left panel. n=9 (Table S3).

The levels of hepatocyte (Cebpa, Hnf4a and Prox1) and biliary epithelial gene expression markers (BEC) (Hnf1b, Epcam, Krt19 and Krt7) in PH and BD organoids were compared with those of freshly isolated primary cells. While the expression of hepatocyte markers was increased in BD organoids following differentiation, the mRNA levels of these markers remained significantly lower than those of freshly isolated hepatocytes (Figure 3). Furthermore, BD organoids increased the expression of BEC lineage markers following differentiation, indicating that the differentiation conditions defined by Huch et al (2013) were permissive for the maturation of the cells towards both hepatocyte and BEC lineages (Figure 3). By contrast, differentiated PH organoids expressed hepatocyte lineage genes at comparable levels with freshly isolated hepatocytes and lacked expression of BEC markers (Figure 3). Altogether these results indicate that differentiated PH organoids maintain a ‘core hepatocyte’ program and respond to Wnt activation as hepatocytes in a resting state; Thus, differentiated PH organoids may be used as a cell source to synthetically reconstruct Wnt-driven zonation.

**Figure 3.**
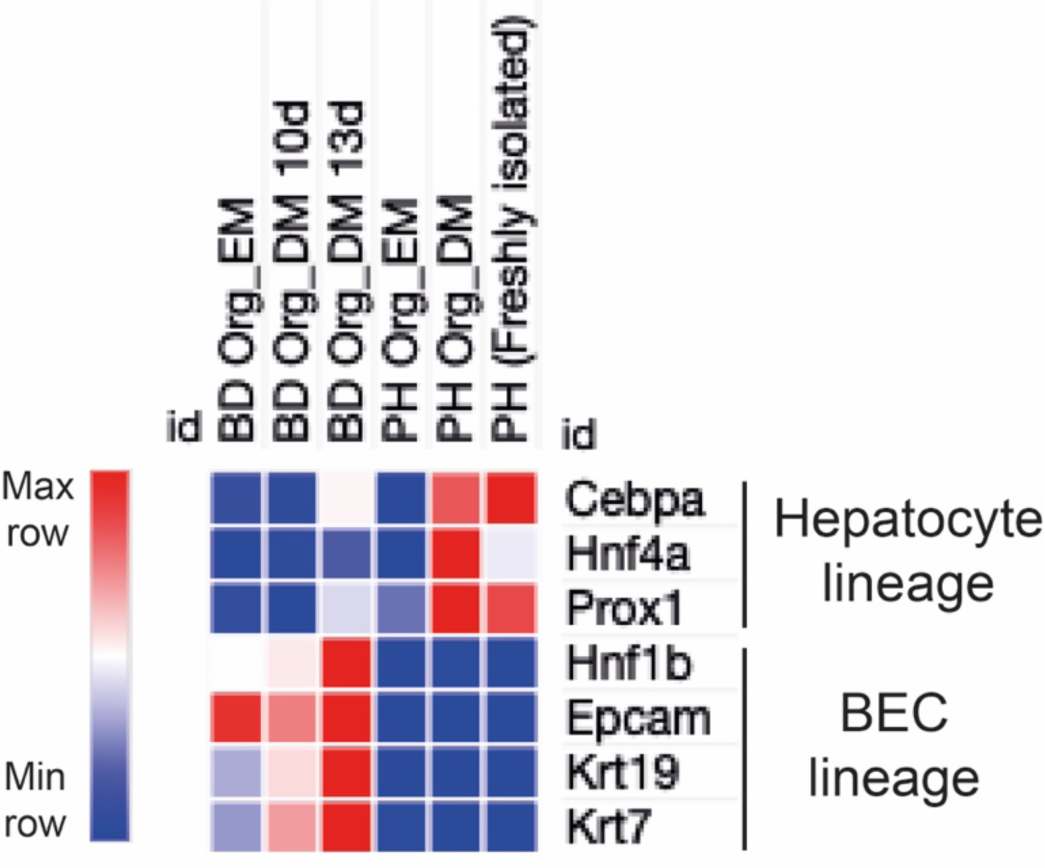
Heat map shows RT-qPCR gene expression levels in BD organoids in expansion medium (BD Org_EM), BD organoids in differentiation medium day 10 (BD Org_DM10d) and in differentiation medium day 13 (BD Org_DM13d); PH organoids in expansion medium (PH Org_EM), PH organoids in differentiation medium (PH Org_DM) and freshly isolated primary hepatocytes (PH). PH organoids were differentiated in the absence of a zonal inducer (Neutral differentiation). Time in culture in expansion medium in BD and PH organoids was matched with their respective differentiation medium conditions. For BD organoid samples, n=6 (Table S3). For PH organoid samples, n=4 (Table S3). For freshly isolated PH samples, n=3 animals.

### Wnt9b and Rspo3 are a superior pericentral inducers to GSK3 inhibitor-based protocols

To investigate how pericentral maturation in the presence of Wnt9b and Rspo3 compared to the original CHIR-based protocol published by Peng et al. (2018), we examined the transcriptome profiles of PH organoids differentiated in the presence of Wnt9b and Rspo3 alone or in combination (W9, R3 and W9R3) and the GSK-3 inhibitor CHIR99021 (CHIR) by RNAseq (Figure 4, A and B). As a control, organoids were differentiated in the absence of zonal inducers (‘neutral’ differentiation) (Figure 4, A and B). It was expected that W9R3 and CHIR, by activating canonical Wnt signaling, would induce similar pericentral maturation programs. When PH organoid differentiation was carried out in the presence of CHIR, 323 transcripts were differentially expressed (up or down regulated) between CHIR and neutral-differentiated organoids (Figure 4, C). However, only 52 transcripts were altered (up or down regulated) following W9R3-treatment (padj <0.05) (Figure 4, C). Unexpectedly, only 15 transcripts (Csad, Cyp2c55, Cyp1a2, Cml2, Ugt2b35, Pcp4l1, Gm22681, Lst1, Gm25803, Wipf3, Ighd3-1, Gm24991, Gm26152, Gm26141 and Gm25670) were significantly altered by both CHIR- and W9R3-treatments when compared to neutral differentiation (Figure 4, C). Of the overlapping set of 15 transcripts, all of them were upregulated with the exception of Csad, that was downregulated only in CHIR-differentiated organoids when compared to ‘neutral’ differentiation (Figure 4, C and Figure 5 A). A table with the differential expression values and significances in each of the treatments when compared to neutral differentiation can be found in Table S1.

**Figure 4.**
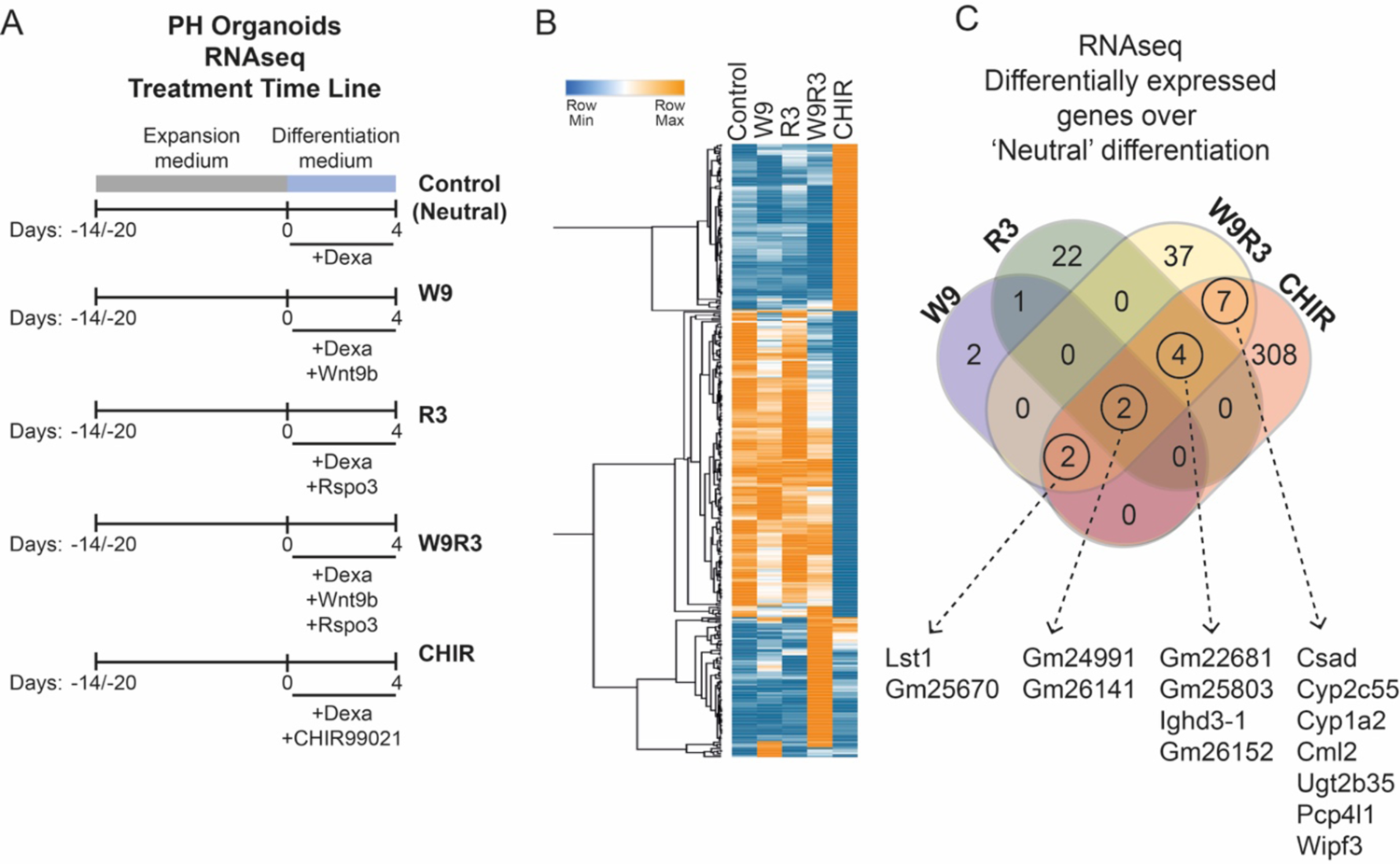
(A) PH organoids differentiation and treatment timeline for RNAseq experiments (n=4, see Table S3). (B) Heatmap showing hierarchical clustering (one minus Pearson correlation) of average RNAseq gene expression values (in normalized CPM) of PH organoids differentiated in the presence of various Wnt pathway activators. Analysis was performed in transcripts differentially expressed (padj value *≤*0.05) between W9-, W9R3-, R3- or CHIR-treated and control organoids. A total of 385 transcripts were found differentially expressed. (C) Diagram shows the number of transcripts differentially expressed (padj value *≤*0.05) in W9-, W9R3-, R3- or CHIR-treated when compared to control organoids. Genes significantly altered by both W9R3 and CHIR treatments when compared to ‘neutral’ differentiation conditions have been highlighted.

**Figure 5.**
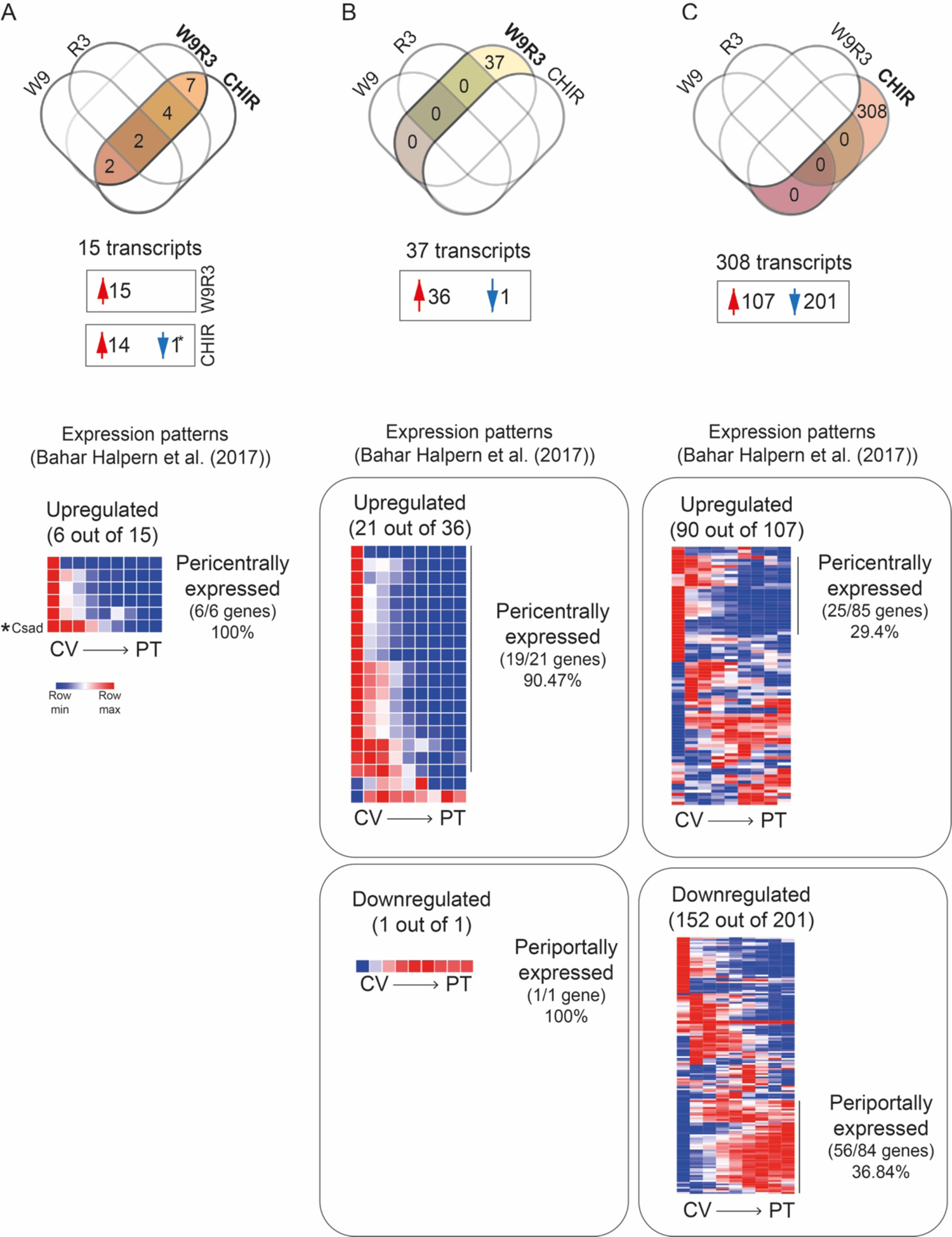
(A) Two top diagrams show the number of transcripts (a total of 15) that were genes differentially expressed (padj value *≤*0.05) in W9R3- and CHIR-treated when compared to control organoids. On the bottom, heatmap shows the patterns of expressions in the central vein (CV)/ portal triad (PT) axis of these transcripts according to Halpern et al. (2018) public data ^8^. The asterisk highlights the row corresponding to the expressions patterns of Csad, the transcript that was downregulated in CHIR conditions. (B) Two top diagrams show the number of transcripts (a total of 37) that were genes differentially expressed (padj value *≤*0.05) in W9R3- but not in CHIR-treated organoids when compared to control organoids. Only one gene appeared significantly downregulated when comparing W9R3-treated and control PH organoids. On the bottom, heatmaps showing the patterns of expressions in the central vein (CV)/ portal triad (PT) axis of these transcripts according to Halpern et al. (2018) public data ^8^. (F) Two top diagrams show the number of transcripts (a total of 308) that were genes differentially expressed (padj value *≤*0.05) in CHIR- but not in W9R3-treated organoids when compared to control organoids. On the bottom, heatmap showing the patterns of expressions in the central vein (CV)/ portal triad (PT) axis of these transcripts according to Halpern et al. (2018) public data ^8^.

To better understand the overlap between the differentially-expressed genes and zonation, we compared the gene sets to the zonal expression profiles that were resolved with a 9-layer resolution by Bahar Halpern et al. (2017) ^8^. It was expected that Wnt-activating treatments would promote the expression of genes expressed pericentrally while repressing the expression of genes that are preferentially expressed by periportal hepatocytes. Most but not all the transcripts detected in our RNAseq analysis were present in the Bahar Halpern et al. (2017) data set. For example, out of the 15 transcripts shared between W9R3 and CHIR only 6 (Csad, Cyp2c55, Cyp1a2, Ugt2b35, Pcp4l1, Wipf3) were present in this data set and therefore only the zonation profiles of these genes could be uncovered (Figure 5, A). These 6 transcripts followed a well-defined pericentral zonation (Figure 5, A). This included Csad, the gene that was downregulated following CHIR differentiation (Figure 5, A). Out of the 36 transcripts upregulated following W9R3 differentiation, only 19 were present in the data set of Bahar Halpern et al. (2017) and 90.47% of these (19 out of 21) were expressed pericentrally (Figure5, B). This percentage contrasted with the low proportion of CHIR-induced genes that were preferentially expressed by pericentral hepatocytes, with only 29.4% of these (25 out of 85 present in Bahar Halpern et al (2017) data set) preferentially expressed by pericentral hepatocytes (Figure5, C). Only one transcript (gene Agt) was downregulated following W9R3 differentiation and, as expected, the expression of this gene was lower in pericentral hepatocytes than in periportal hepatocytes (Figure5, B). This again contrasted with the patterns of expression of genes downregulated following CHIR-differentiation, with only 36.84% of them (152 out of 201) enriched in periportal hepatocytes (Figure 5, C). Altogether these results suggests that a combination of Wnt9b and Rspo3 induce a more specific pericentral maturation than the original CHIR-base protocol.

We next investigated the induction of well-established β-catenin target genes in PH organoids following differentiation in the presence of the different Wnt pathway activators. As expected, RNAseq analysis indicated that differentiation in the presence of W9R3 but not with the individual ligands (W9 and R3) caused a significant increase in Axin2 mRNA levels and also induced the expression of other Wnt target genes including Rnase4, Notum, Lect2 and Tbx3 (Figure 6, A). Lgr5 and Lgr4, two membrane receptors mediating Rspo3 signaling, were identified as upstream regulators by Ingenuity Pathway Analysis (IPA) in W9R3-differentiated organoids (Figure 6, D). Surprisingly, none of these genes, including Axin2, were significantly induced following differentiation in the presence of CHIR at the timepoints and concentrations used (Figure 6, A). Principal component analysis revealed that the sequenced PH organoids preferentially clustered by ‘experimental set’ between the four experimental rather than by treatment (Figure S1), suggesting that inter-sample ‘noise’ within the datasets may obscure more subtle variations in gene expression that may overlap with inter-sample biological differences.

**Figure 6.**
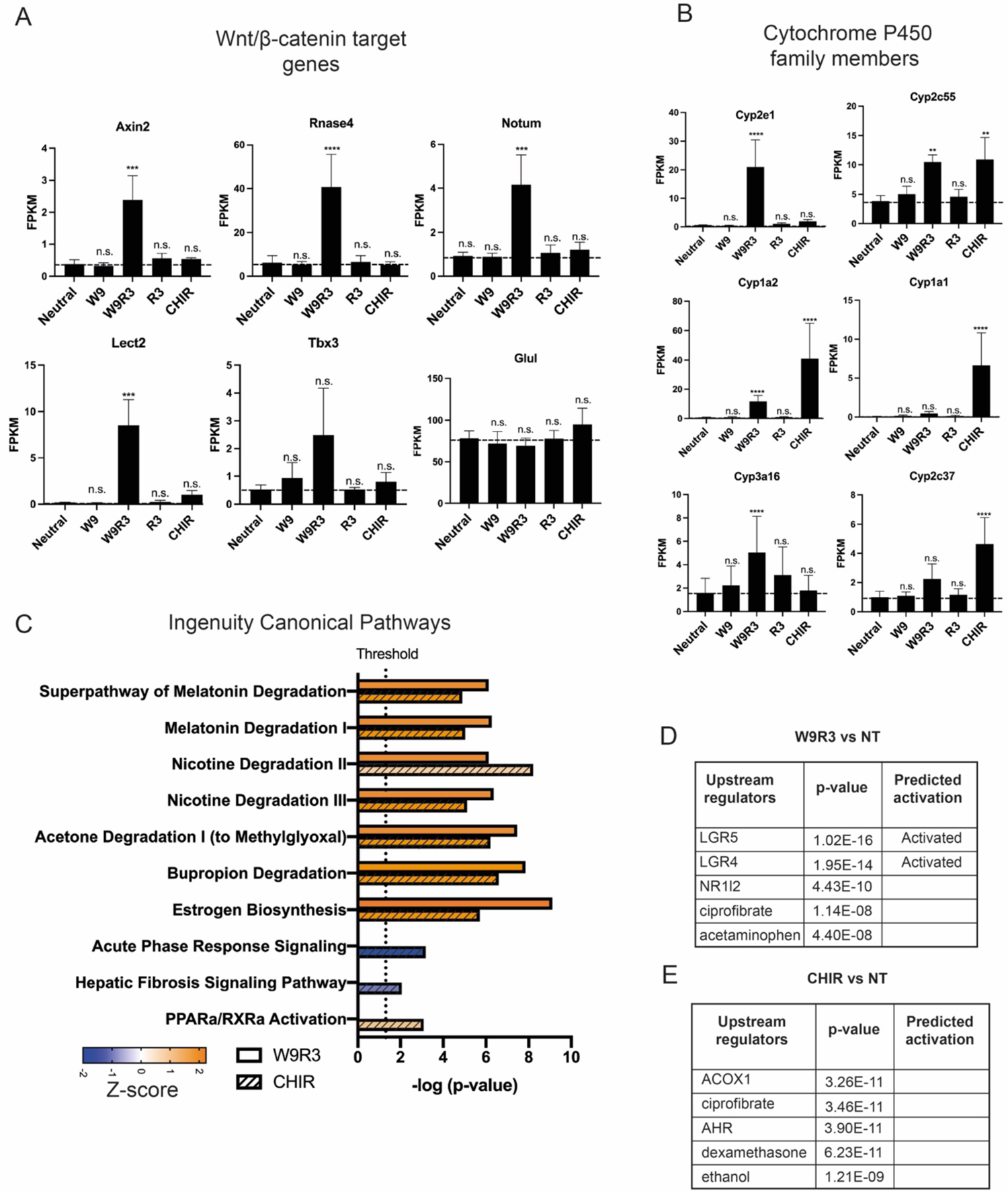
(A and B) Expression levels of well-established Wnt/*β*-catenin target genes and members of the cytochrome P450 family in PH organoids differentiated in the absence of a Wnt inducer (control) or in the presence of W9, W9R3, R3 or CHIR by RNAseq expressed in normalized FPKM values. (n=4, see Table S3). (C-E) IPA canonical pathway analysis (panel C) and IPA upstream regulators analysis (panel D and E) of differentially expressed genes (padj value <0.05) between W9R3-treated and control PH organoids (a total of 52 transcripts) or CHIR-treated and control PH organoids (a total of 323 transcripts).

β-catenin induces the expression of members of the cytochrome P450 family in pericentral hepatocytes. Subsets of cytochrome P450 enzymes were differentially regulated by W9R3 or CHIR. Cyp1a2 and Cyp2c37 were induced by both W9R3 and CHIR. Cyp2e1 and Cyp3a16 were induced by W9R3 but not CHIR (Figure 6, B). Conversely, Cyp1a1 and Cyp2c37 were induced by CHIR but not W9R3 (Figure 6, B). Top IPA canonical pathways enriched in W9R3- and CHIR-differentiated organoids were the ‘Superpathway of Melatonin Degradation’, ‘Nicotine degradation II and II’, ‘Acetone Degradation I to Methylglyoxal’ and ‘Buprion degradation’ (Figure 6, C). These probably reflect the activation of members of the cytochrome P450 family by both W9R3 and CHIR treatments. Following treatment with CHIR, the activation of centrally-expressed cytochrome P450 enzymes by CHIR may similarly be related to the induction of zonal liver markers. However, it cannot be excluded that some changes are related to the metabolic processing of the CHIR compound itself, although further experiments are require to determine whether this is the case.

Gene Set Enrichment Analysis (GSEA) was used to compare organoids differentiated in the presence of CHIR- or W9R3. CHIR-differentiated organoids were positively enriched in gene sets previously associated with stem cells and with Sox9 transcriptional activity in embryonic stem cells (Figure 7 A and B). CHIR-differentiated organoids were also enriched in genes associated with RASD2, a small GTPase belonging to the Ras superfamily (Figure 7, C). Although the link between RASD2 and CHIR’s target is unclear, GSK3β is a kinase that also coordinates the activation of multiple signalling cascades including phosphoinositide 3-kinase, Notch, and Hedgehog signalling pathways ^37^. Two independent groups have shown that CHIR is critical for the sustained proliferation and long-term culture of PH organoids in the expansion phase ^33, 34^. Interestingly, Lv et al. (2014) found that the canonical ligand Wnt3a could not replace the need of CHIR to support primary hepatoblast self-renewal in culture ^38^. Taken together, this suggests that the broad inhibition of GSK-3 activity may be required for the expansion of PH organoids in a stem-like state, but that during differentiation towards a peri-central state, a specific combination of Wnt pathway ligands is required.

**Figure 7.**
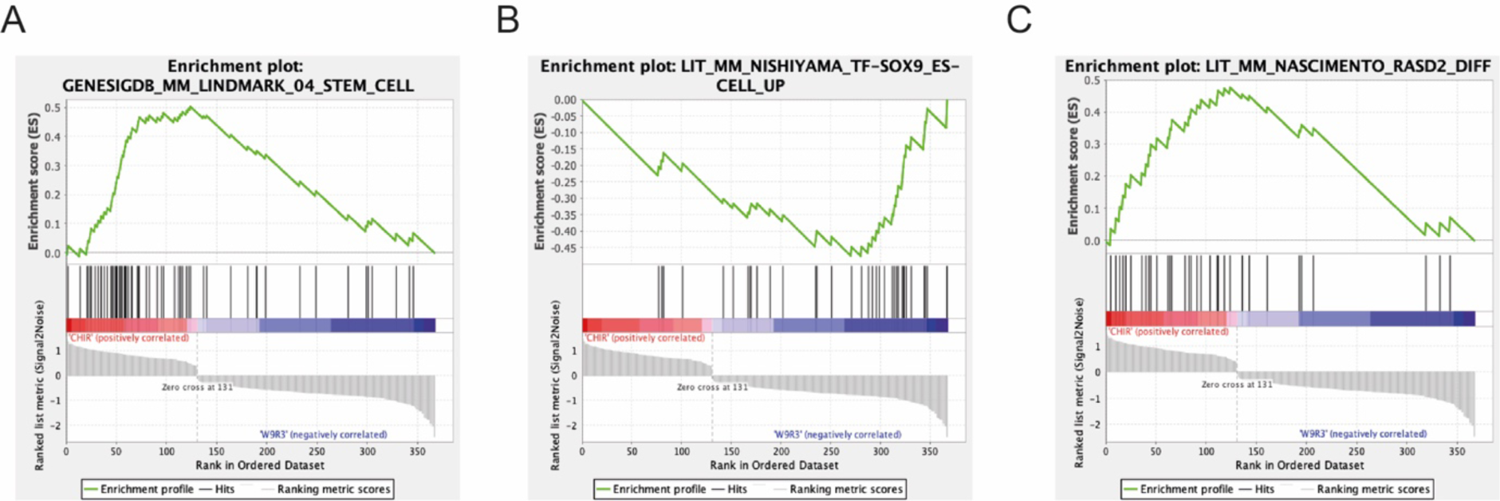
GSEA analysis using C2 comprising Curated Gene Sets from online pathway databases publications in Pubmed and knowledge of domain experts reveal enrichment of stem cell related genes (panel A) in CHIR-differentiated PH organoids when compared to W9R3-differentiated PH organoids. W9R3-differentiated organoids were negatively enriched for Sox9 stem cell related genes when compared to CHIR-treated organoids (panel B). CHIR-differentiated organoids were enriched in RasD2-related genes (panel B). Analysis was performed in genes differentially expressed genes between CHIR- and W9R3-treated organoids with a padj < 0.05. A total of 490 genes where found differentially expressed. Of those 430 had a human orthologue and therefore could be analyzed using GSEA.

Peng et al. (2018) reported that differentiation in the presence of CHIR or EGF/HGF increased the expression of members of hepatocyte pericentral and periportal genes, respectively, when compared to expansion conditions ^33^. However this study did not carry out a side-by-side comparison of CHIR- and EGF/HGF-based differentiation conditions. As the data presented here raise the possibility that CHIR can function as both an inducer of both a stem cell program and an inducer pericentral zone gene expression, we repeated Peng et al. (2018)’s study but also compared the expression of pericentral and periportal genes in EGF/HGF- and CHIR-differentiated PH organoids with ‘neutral’ differentiation conditions) (Figure 8, A). As previously reported, EGF/HGF-differentiated organoids expressed higher levels of periportal markers (Alb, Ass1 and Cyp2f2) when compared to PHorg-EM organoids (Figure 8, B) ^33^. Similarly, differentiation of organoids in the presence of CHIR significantly induced the expression of the pericentral genes Cyp1a2, Cyp2e1 and Fah when compared with expansion conditions (PHorg-EM) (Figure 8, B) ^33^. However, the expression levels of the pericentral genes Cyp1a2, Cyp2e1 and Fah remained at comparable levels between in CHIR- and ‘neutral’-differentiated organoids (Figure 8, C). Similarly, and with the exception of Alb, periportal genes were also expressed at comparable levels between in EGF/HGF- and ‘neutral’-differentiated organoids (Figure 8, B and C). In summary, while the data published by Peng et al (2018) was replicated, both pericentral and periportal genes expression were ‘induced’ in neutral differentiation conditions ^33^. This suggests that much of the increase in periportal and pericentral markers expression reported by Peng et al. (2018) was driven by the addition of dexamethasone and not by the EGF/HGF or CHIR zonal inducers.

**Figure 8.**
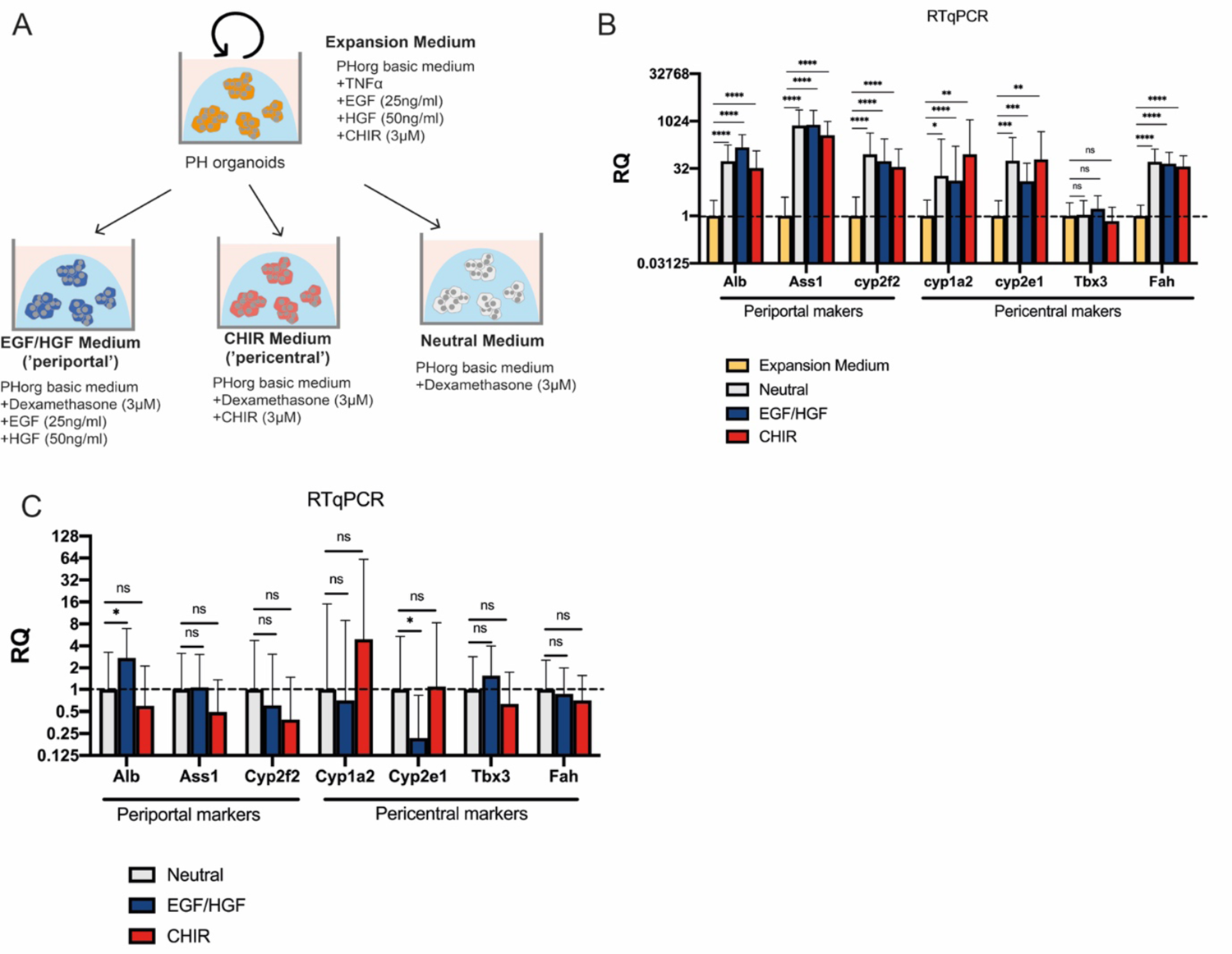
(A) PH organoid differentiation scheme where the components defining each of the media have been highlighted. (B) RT-qPCR gene expression analysis showing the mRNA levels of genes preferentially expressed by pericentral or periportal hepatocytes normalized over the expression values of PH organoids cultured in expansion conditions. B2M was used as a house keeping gene. (n=5). RT-qPCR gene expression analysis where mRNA expression levels of were normalized over the expression values of PH organoids differentiated in neutral conditions. B2M was used as a house keeping gene. (n=5). Mann-Whitney t-test (p-value n.s.>0.05; p-value *< 0.05; p-value **< 0.01; p-value ***< 0.001; p-value ****< 0.0001).

### Immobilized Rspo3 in combination with soluble Wnt9b locally activates the canonical in hepatocyte and HEK293 cell cultures

Having determined that Wnt9b and Rspo3 proteins induce a pericentral metabolic program in PH organoids, we next sought to determine a strategy for the local presentation of these cues *in vitro*. Wnt9b and Rspo3 proteins were immobilized on EDC-treated carboxylic acid-coated beads which react with free amine groups on the proteins to form zero linker length conjugates (Figure S2) ^39^. Activated beads that had been quenched with BSA alone were used as a control.

The activity of soluble and immobilized Wnt9b and Rspo3 ligands was assessed in HEK293T cells stably expressing a 7xTCF luciferase (TCF-Luc) reporter ^40^. Wnt9b only induced TCF-dependent transcription at any of the concentrations tested (20 ng/ml to 500 ng/ml), when it was added the presence of Rspo3 (10 ng/ml) (Figure 9A-C). These data contrasted with the Rspo-independent activation of TCF-dependent transcription by Wnt3a (Figure S3) and are similar to previous reports using murine hepatocyte and kidney cells ^20, 41^. Soluble Rspo3 protein alone did induce a low level of TCF-dependent transcription alone, but this was over 15-fold lower than that observed in the presence of both ligands (Figure 9, C). The induction of the TCF-Luc reporter by Rspo3 was abrogated when the reporter cells were first treated with LGK974, a porcupine inhibitor that blocks the secretion of Wnt proteins (Figure 9, D), suggesting that Rspo-3 induction of TCF-dependent transcription was dependent on the synthesis and secretion of endogenous Wnt ligands. This suggestion was further supported by the observation that LGK974 inhibition could be reversed by the addition of exogenous Wnt9b soluble protein in the presence of Rspo3 (Figure 9 D).

**Figure 9.**
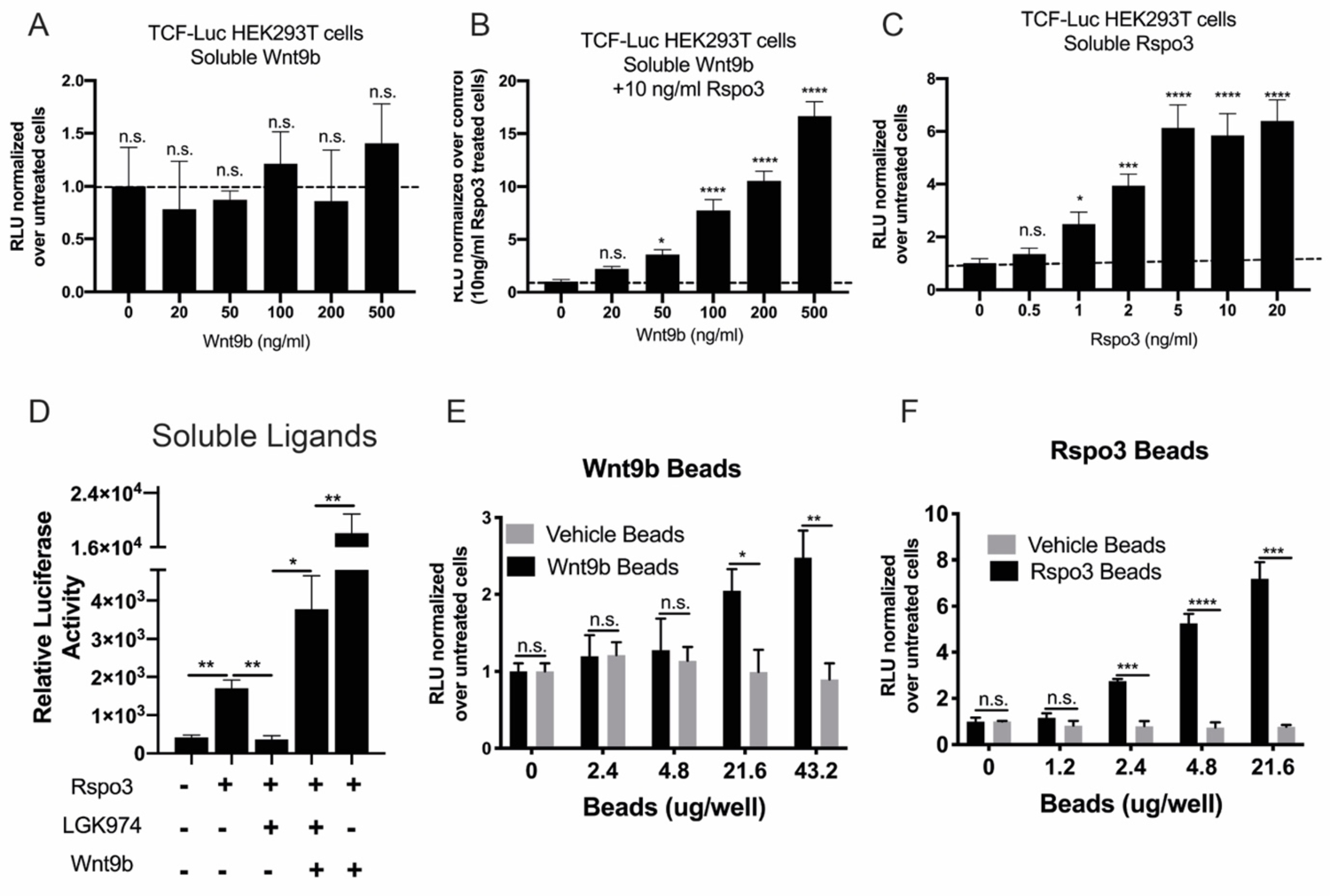
(A-C) 24h response of HEK293TTCF-Luc reporter cells to different concentrations of soluble Wnt9b and Rspo3 ligands. (D) Blockage of endogenous Wnt secretion (exposure to 500 nM LGK974) depletes Rspo3 (10 ng/ml) mediated activation of the TCF reporter. Addition of Wnt9b (100 ng/ml) is sufficient to activate the reporter.in the presence of Rspo3. LGK974 was administrated 24h prior to Wnt9b or Rspo3 exposure and maintained during the whole length of the experiment. (E-F) Response of HEK293T TCF-Luc reporter cells to the exposure of immobilized Wnt and Rspo ligands for 24h. Wnt9b and Rspo3 beads but not vehicle beads triggers the activation of the TCF-Luc reporter in a dose dependent manner. Carboxylic acid beads activated with EDC/NHS and quenched in 0.1% BSA PBS were used as a vehicle beads in all the experiments. Mann-Whitney t-test (p-value n.s.>0.05; p-value *< 0.05; p-value **< 0.01; p-value ***< 0.001; p-value ****< 0.0001).

The activity of Wnt9b and Rspo3 conjugated beads was assessed in the TCF-luciferase reporter cells. Wnt9b beads when added in the presence of soluble Rspo3 (10ng/ml) induced TCF dependent transcription in a bead-concentration dependent manner (Fig. 9, E). Similarly, Rspo3-beads activated TCF-dependent transcription in a concentration dependent manner (Compare Figure 9 C and F). Given that Rspo3 beads induced the reporter more robustly than Wnt9b beads (Compare Figure 9 E and F), subsequent studies focused on the activity of immobilized Rspo3 with and without soluble Wnt9b. To allow localized activation to be assessed, a TCF-eGFP monoclonal cell line was generated by transducing HEK293T cells with lentivirus vectors harboring the 7xTCF eGFP reporter. Clones that were highly sensitive to Wnt/β-catenin activation were isolated by FACs on the basis of eGFP expression in the presence of Wnt3a (100ng/ml and 200ng/ml) treatment, a well-established canonical Wnt ligand (Figure S4). Both soluble and immobilized Rspo3 protein induced the appearance of eGFP positive cells in a dose dependent manner by flow cytometry analysis (Figure 10, A and B). The local activity of Rspo3 beads was next evaluated in HEK293T eGFP cells by fluorescence microscopy. Movement of the Rspo3 beads in the cultures was prevented by embedding the beads into collagen (Figure 10, C). Collagen-embedded beads were brought into close proximity to the HEK293T cultures using magnetic fields generated with a DynaMag-2 magnet prior to collagen jellification. The pulling of Rspo3 beads also caused the aggregation of the beads into clusters (Figure 10, D). Presentation of Rspo3 beads in combination with soluble Wnt9b caused the local activation of the Wnt/β-catenin pathway in HEK2913 cultures, suggesting that a combination of soluble Wnt9b and localized Rspo3 might be an attractive way to recreate the local Wnt/Rspo microenvironment of the central vein (Figure 10, E).

**Figure 10.**
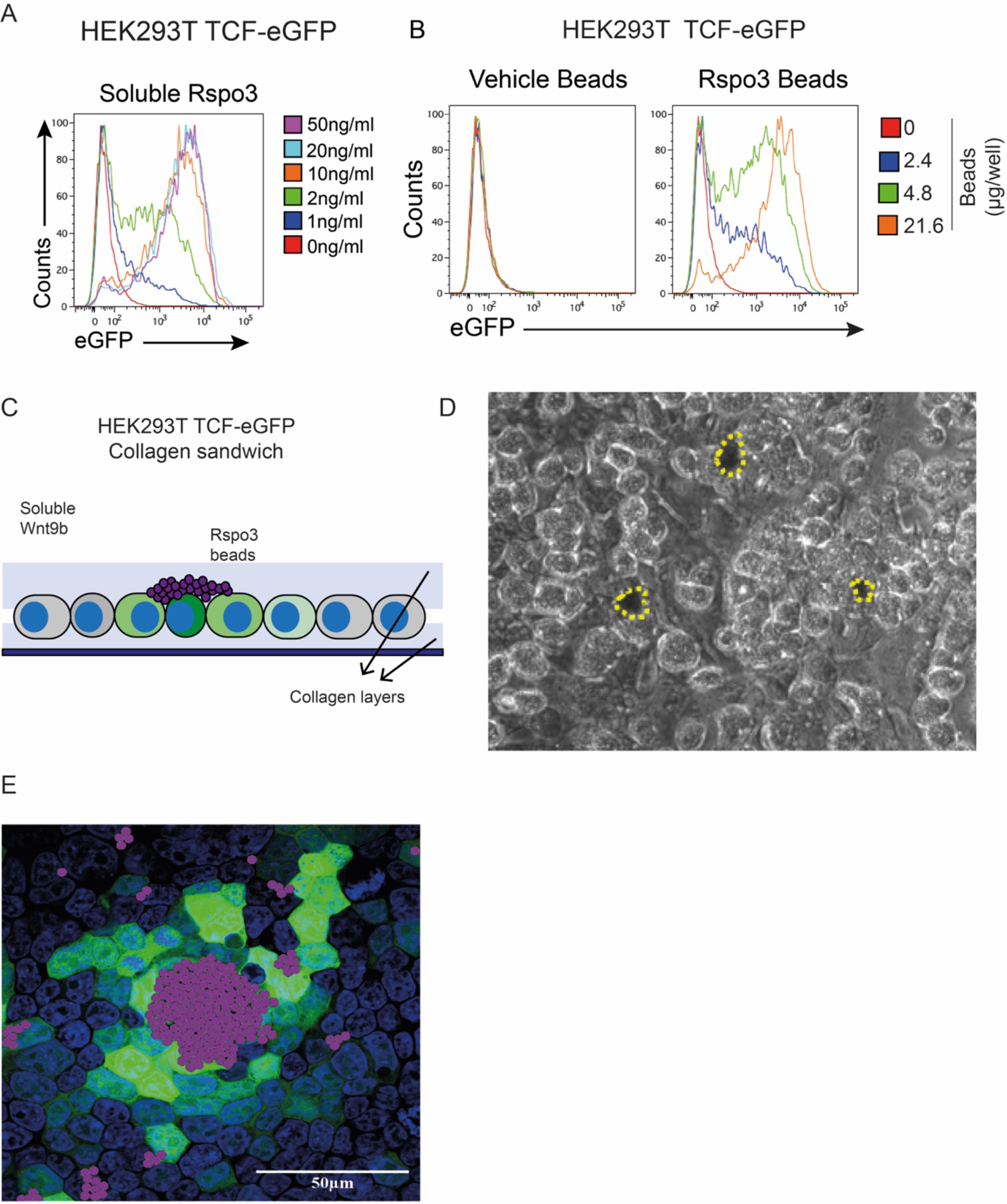
(A) Flow cytometry plots showing 24h dose response of HEK293T TCF-eGFP cells (clone #47) to soluble Rspo3. HEK293T TCF-eGFP cells are responsive to Rspo3 in absence of exogenous Wnt ligands. (B) Flow cytometry histograms showing eGFP fluorescence levels in HEK293T TCF-eGFP reporter cells after the exposure to Wnt/Rspo immobilized ligands. Rspo3 beads induce the expression of eGFP in a dose dependent manner whereas the TCF-eGFP reporter remains unaffected by the presence of vehicle beads (beads activated with EDC/NHS and quenched in 0.1% BSA PBS). (C) Collagen sandwich culture strategy diagram. (D) Image shows clustering of beads (yellow dash line) in HEK293T cells cultured in a collagen sandwich conformation. (D) HEK293T TCF-eGFP cells cultured in a collagen sandwich platform were exposed to Rspo3 beads and soluble Wnt9b for 24h. Beads were artificially coloured in purple.

The synthetic reconstruction of zonation will ultimately require the simultaneous incorporation of contra-opposed Wnt and oxygen gradients; thus, strategies to induce Wnt-driven zonation must achieve local activation of the Wnt/*β*-catenin cascade in a controlled manner and be compatible with flow culture systems that introduce oxygen and metabolite gradients. In the present study we showed that Wnt9b and Rspo3 proteins can be conjugated onto iron beads using a two-step EDC/NHS immobilization procedure and therefore provide a proof-of-principle that immobilization of Rspo3 in combination with soluble Wnt9b may be a valid strategy to artificially reconstruct the Wnt niche of the hepatic central vein in vitro. To be able to fully investigate interactions between immobilized Wnt pathway ligands and PH organoid-derived hepatocytes in will require the generation of novel strategies to engineer intimate interactions between cells and synthetic surfaces. This work lies outside the scope of this investigation, but would be expected to generate systems of great interest for those interested in modelling normal liver function for studies of toxicology and liver metabolism. Approaches could involve the seeding of PH organoids within collagen sandwiches as previously described or the more ambitious integration of PH organoid derived cells within 3D structures incorporating matrigel. The tools developed within this report should strongly support these developments (^42–44^.

## Methods

### Isolation and culture of primary hepatocytes

PHs were isolated following a classical two-step collagenase perfusion method with modifications using two aquarium pumps (EHEIM, Typ 1000.340, Serie 14042) connected to a home-made silicone tubing system. Briefly, murine livers were perfused through the portal vein at a 5-10 ml/min first with Solution I (EBSS without calcium, magnesium and phenol red, supplemented with 10 mM HEPES and 0.5 mM EGTA) for 8-15min and then with Solution II (DMEM/F12 no phenol red supplemented with 10 mM HEPES and 0.6-0.8 mg/ml collagenase type I) until signs of digestion were apparent. Both solutions I and II were pre-warmed at 39C prior to perfusion. After digestion, primary hepatocytes were released from the hepatic sac into hepatocyte wash medium (DMEM/F12 no phenol red (Invitrogen) supplemented with 10 mM HEPES (Invitrogen), 1:1000 ITS and 1% Pen Strep) and purified by three slow centrifugations (3x 3min at 30g). Cell viability of the last hepatocyte enriched fraction was determined by acridine orange (AO)/ propidium iodide (PI) dual staining (Thermofisher) using a LUNA cell counting system (LabTech). Only hepatocytes from isolations with a viability >70% were used in this study. Hepatocytes in hepatocyte attachment medium (1:1000 ITS, 5% ES qualified FBS, 1% Pen/Strep, 15 mM HEPES DMEM/F12 no phenol red) were seeded onto plates coated with 50 μg/ml collagen type I from rat tail (ThermoFisher) for 1 to 2h at room temperature. After 16h attachment, hepatocytes were washed with pre-warmed PBS and medium was replaced for hepatocyte culture medium (1:1000 ITS, 0.5% ES qualified FBS, 1% Pen/Strep, 15 mM HEPES DMEM/F12 no phenol red) supplemented with Wnt9b (100 ng/ml) or Rspo3 (50 ng/ml) when indicated. Medium was replaced every day.

### Generation and culture of PH organoids

PH organoids were generated from primary hepatocytes as previously described ^33^. Briefly, primary hepatocytes isolated as before were resuspended in ice-cold Matrigel at a concentration of 50 to 100 cells/μl in 10-20x 25μl hydrogel drops on 6-well plates pre-warmed at 37°C. A maximum 1:5 ratio of hepatocyte suspension:Matrigel was used. After 14 to 20 days, organoids reached an average size of 300-1000 μm and were ready for passage. For organoid passage, organoids were washed 1x in ice-cold PBS and treated with TrypLE (Gibco) for 5-10min with gentle re-suspension using a p200 every 3min. Upon organoid breakage onto clusters of 5 to 10 cells, TrypLE digestion was interrupted by adding ice-cold William’s E media and organoids pelleted (300 g, 5 min) and resuspended in Matrigel. Organoids were routinely maintained at a split ratio of 1:2 every 14 to 20 days. During expansion, organoids were cultured in PHorg-EM (Williams E media (no phenol red) supplemented with 1% GlutaMAX, 10 nM HEPES, 1% Pen/Strep, 2% B27 supplement, 1% N2 supplement, 10 mM Nicotinamide, 1.25 mM N-acetylcysteine, 10 μM Y-27632, 1 μM A83-01, 1% (v/v) Non-essential amino acids, 3 μM CHIR99021, 25 ng/ml EGF, 50ng/ml HGF, 100ng/ml TNF*α*). From passage 2 onwards, PHorg-EM was supplemented with 50 ng/ml Noggin for the first 7 days of culture. For differentiation, organoids were switched to PHorg-DM (Williams E media (no phenol red) supplemented with 1% GlutaMAX, 10 nM HEPES, 1% Pen/Strep, 2% B27 supplement, 1% N2 supplement, 10 mM Nicotinamide, 1.25 mM N-acetylcysteine, 10 μM Y-27632, 1 μM A83-01, 1% (v/v) Non-essential amino acids, 3 μM dexamethasone) supplemented with Wnt9b (100 ng/ml), Rspo3 (50 ng/ml), CHIR99021 (3 μM) or EGF (25 ng/ml) and HGF (50ng/ml) when indicated. PH organoids used in this study were between passage 5 and 10.

### Generation and culture of BD organoids

BD organoids were generated from hand-picked bile ducts as previously described ^32^. BD organoid expansion medium was defined as AdDMEM/F12 containing 1% B27 supplement (Invitrogen), 0.5% N2 supplement (Invitrogen), 1.25 uM N-Acetylcysteine (Sigma), 10 nM gastrin (Sigma),100 ng/ml FGF10 (Prepotech), 50ng/ml EGF (Peprotech), 50 ng/ml HGF (Peprotech), 10 nM Nicotinamide (Sigma) and 10% Rspo1 conditioned medium prepared in house. For the first 7 days of culture after isolation, expansion medium was supplemented with 25 ng/ml Noggin (Prepotech). Organoids were routinely 1:5 passaged by mechanical disruption every 5 to 8 days. When seeded for experiment, organoids were incubated with TriplE (Gibco) for 7 min at 37°C, digested into single cells and seeded at a concentration of 200-400 cells/μl matrigel. For differentiation, organoids were expanded for 2 to 4 days, and switched to differentiation media (AdDMEM/F12 containing 1% B27 supplement (Invitrogen), 0.5% N2 supplement (Invitrogen), 1.25 μM N-Acetylcysteine (Sigma), 50 ng/ml EGF (Peprotech), 100 ng/ml FGF10 (Peprotech), 50nM A-8301 (Prepotech), 10 μM DAPT (Sigma)) and cultured for 13 days. Differentiation media was changed every day and supplemented with 3 μM dexamethasone for the last 3 days of culture. When indicated, differentiated organoids were treated with 100 ng/ml Rspo1 in 0.1% BSA PBS or Doxycyclin (2 mg/ml in PBS). BD organoids used in this study were between passage 4 and 15.

### RNA extraction and RT-qPCR

RNA extraction was performed by using TRIzol® (life technologies) as indicated by the manufacturer’s instructions with an extra purification step using the RNesy plus kit (Qiagen) of the aqueous phase containing total RNA. cDNA was synthetized using a cDNA synthesis kit (Bio-Rad). Then, cDNA was 1:4 diluted in water and mixed with SYBR® Green (Bio-Rad) and the appropriate primers at 5µM. Three technical replicates were performed per condition. qPCR reaction was carried out on a QuantStudio 7 Flex Real-Time 384-well PCR system (Thermofisher). Quantified expression values for individual genes were normalized to B2M. The gene-specific primer pairs used in this study are in TableS2.

### RNA sequencing

Total RNA from organoids was isolated as previously and RNA concentration was determined by a Hi-sensitivity Qubic kit. Single-end 75bp sequencing was performed with 10ng of total RNA input on a NextSeq-500 sequencer where equi-molar libraries were pooled (Genomic Hub, School of Biosciences, Cardiff University, Cardiff, CF10 3AX, UK). Sequences were trimmed with Trimmomatic 0.35 and assessed for quality using FastQC 0.11.2 ^45^. On average, 99% of the total reads were mapped to the mouse GRCm38 reference genome using STAR 2.5.1b. and counts were assigned to transcripts using featureCounts with the GRCm38.84 Ensembl gene build GTF ^46 47^. Each sample was sequenced twice and the total reads were concatenated after being mapped. Counts per million (CPM) and fragments per kilobase per million (FPKM) were calculated for each gene. Both the reference genome and GTF were downloaded from the Ensembl FTP site (Data Hub, Cardiff University, Cardiff, CF14 4AXN). Genes with <2 average FPKM in all biological groups were considered as not expressed and discarded from the analysis. Differential gene expression was performed using DESeq2 R package, with default 0.1FDR, controlling for biological replicates differences ^48^. The normalised gene expression values (default DESeq2 scaling-factor normalisation) of significantly expressed genes were subject to Principal Component Analysis (PCA) using in R using prcomp(). Genes with adjusted P value (padj) *≤*0.05 were considered as differentially expressed. Gene set enrichment and pathway enrichment analysis was performed using GSEA and IPA, respectively. Heat maps generation and hierarchical clustering with CPM values were performed applying one minus Pearson correlation clustering method using the Broad Institute online tool Morpheus (https://software.broadinstitute.org/morpheus/).

### Immobilization of Wnt9b and Rspo3 proteins onto magnetic beads

Recombinant mouse Wnt9b (carrier free) and Rspo3 (carrier free) was purchased from R&D systems. Wnt9b (25 μg) and Rspo3 (25 μg) were immobilized onto 2.49 mg and 1 mg, respectively, of Dynabeads M-270 Carboxylic acid (Invitrogen) by using a two-step coating procedure with EDC and NHS as indicated by the manufacturer. Briefly, beads were washed twice for 10 min with ice cold 25 mM MES buffer (pH 5) at a concentration of 10 mg beads/ml. Immediately before use, EDC and NHS were respectively dissolved at 50 mg/ml in ice cold 25 mM MES buffer (pH 5). Beads were then incubated at a final concentration of 10 mg/ml in EDC:NHS (1:1) at RT with slow tilt rotation for 30 min and washed 1x with 25 mM MES buffer (pH 5). After this step, the beads were activated and open to form covalent bounds with the primary amines of the ligand. The lyophilized ligands of interest were reconstituted in PBS immediately prior to use and added directly to the activated beads. 25 mM MES buffer (pH 5) was added in 4:5 ratio and the protein/bead mixture was incubated for 1h at RT with slow tilt rotation. Beads containing the immobilized ligands were then washed 3x with PBS and quenched and stored in 1% BSA PBS. As a control, vehicle-beads that have undergo activation and 1% BSA quenching were used. Beads were pulled with a magnet for 4min between washes.

### Generation of a HEK293T TCF-eGFP reporter cell line

p7TGC was a gift from Dr. Roel Nusse (Addgene, #24304). Lentivirus were generated by triple transfection of HEK293T cells with the envelop plasmid VSV-G (3 μg), the packing plasmid psPAX2 (6.5 μg), and the TCF-eGFP reporter plasmid (p7TGC) (10 μg), by calcium-phosphate. Lentivirus were collected 24h post-transfection and stored at −80C for further use. To transduce HEK293T cells with the TCF-eGFP reporter, 50.000 cells/well were seeded into 24-well plates, cultured for 24 h and exposed to different amounts 100 μl of lentivirus-containing medium for 24 h, washed two times with PBS and cultured in 10%FBS DMEM/High glucose for other 24 h. Cells were then collected using Trypsin and cells stably expressing the p7TGC were sorted based on mCherry expression by flow cytometry. To generate a Wnt super-responder monoclonal HEK293T TCF-eGFP reporter cell line, cells stably expressing the p7TGC construct growing at exponential rate were exposed for 24h to 100 ng/ml of Wnt3a and their eGFP protein levels were assessed by flow cytometry. Cells with high eGFP expression were individually sorted into a 96-well plate format. A total of 16 clones were then expanded and their response to 24h exposure to Wnt3a at various concentrations (10, 20, 50 and 100 ng/ml) was evaluated. The most sensitive clone to Wnt3a with the lower eGFP background was used in this study (Figure S4).

### TCF-Luc reporter assay

HEK293T cells stably transduced with the SuperTOPFlash-luciferase reporter were kindly provided by Prof. Nathan (Johns Hopkins University) and cultured in DMEM High Glucose supplemented with 10% FBS and 100 μg/ml of G418 to avoid silencing of the transgene. To measure the response to Wnt ligands, 5000cells/well were plated onto clear-bottom 96-well plates, cultured for two days and then treated with different concentrations and combinations of Wnt ligands as indicated. After 24h, cells were washed with PBS and treated with 50 μl PBS plus 50 μl Blight-Glo (Promega) for 5min in the dark. Luminescence was subsequently measured by a BMG-luminometer.

### TCF-eGFP reporter assay

Cells from the Wnt super-responder monoclonal HEK293T TCF-eGFP reporter line generated as previously indicated (10000 cells/well) were seeded onto 48-well plates, cultured for 2 days and subsequently treated with different concentrations of Rspo3 (soluble or immobilized) in the absence of exogenous Wnt ligands. After 24h treatment, cells were trypsinized and the expression of eGFP was assessed by flow cytometry.

### Clustering of Rspo3 beads in KEH293T sandwich cultures

The solution for the first (1.5 mg/ml) and second (0.9 mg/ml) layer of rat tail collagen type I (Thermofisher) was prepared following the jelling procedure instructions of the manufacturer. 100 μl of collagen layer1 mixture were pipetted per well of an 8-well ibidi chamber slide and distributed using a p200 tip and plates were incubated for 30-45 min at 37°C. After a firm layer1 gel was formed, HEK297T cells were seeded on top of it. 12-16h later, dead cells and non-attached cells were washed one time with 37°C pre-warmed PBS, the second layer of collagen with the beads embedded was placed on top of them. Beads were attracted to the bottom of the plates by placing the plates on top of a DynaMag-2 magnet for 2-3 second and the top layer of collagen was then jellified for 30-45min at 37°C. After the second collagen layer was solidified, culture medium was pipetted on top.

## Statistical analysis

GraphPad Prism 7.0 Software was used for all statistical analysis and graphs representation. Statistics for RT-qPCR data were determined using Mann-Whitney test. SD error bars. Heat-maps for RT-qPCR data representation were generated with the online tool Morpheus (www.software.broadinstitute.org/morpheus/).

## Conflict of Interest

The authors declare no competing interests.

## Acknowledgments

The TCF-Luc HEK293T cell line was a gift of Dr Jeremy Nathans (Johns Hopkins University). We thank Dr Matt J Smalley from Cardiff University for helpful discussions. We thank Angela M. Marchbank and Georgina E. Smethurst for the RNAseq library preparation and Robert Andrews and Sumukh Deshpande for assistance with the bioinformatic analysis. This work was supported by the Marie Sklodowska-Curie Actions Innovative Training Network 675407 (acronym PolarNet) and the William Morgan Thomas Fund. M.M., and T.G.B. were funded by the Wellcome Trust (Grant number: WT107492Z).

## Supporting Information

**Figure S1.**
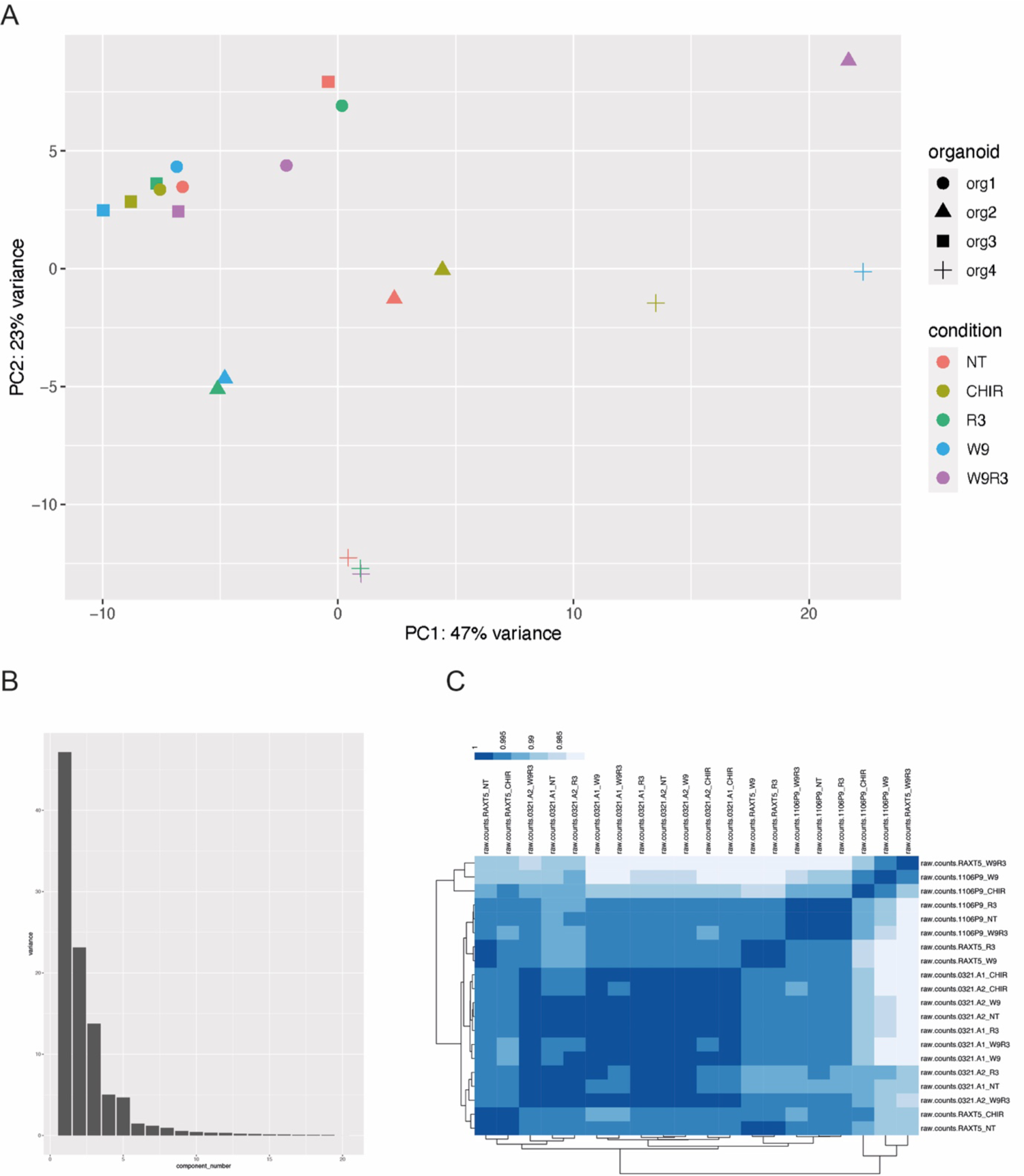
(A) PCA of PH organoids differentiated in Neutral (NT), CHIR, R3, W9 or W9R3 differentiation conditions shows preferential clustering of the samples based on the animal of origin or experimental set rather than by treatments. (B) Scree plot showing the eigenvalues for each of the dimensions of the PCA. (C) Heatmap showing hierarchical clustering of PH organoids differentiated in Neutral (NT), CHIR, R3, W9 or W9R3 differentiation conditions based on sample distances calculated with R.

**Figure S2.**
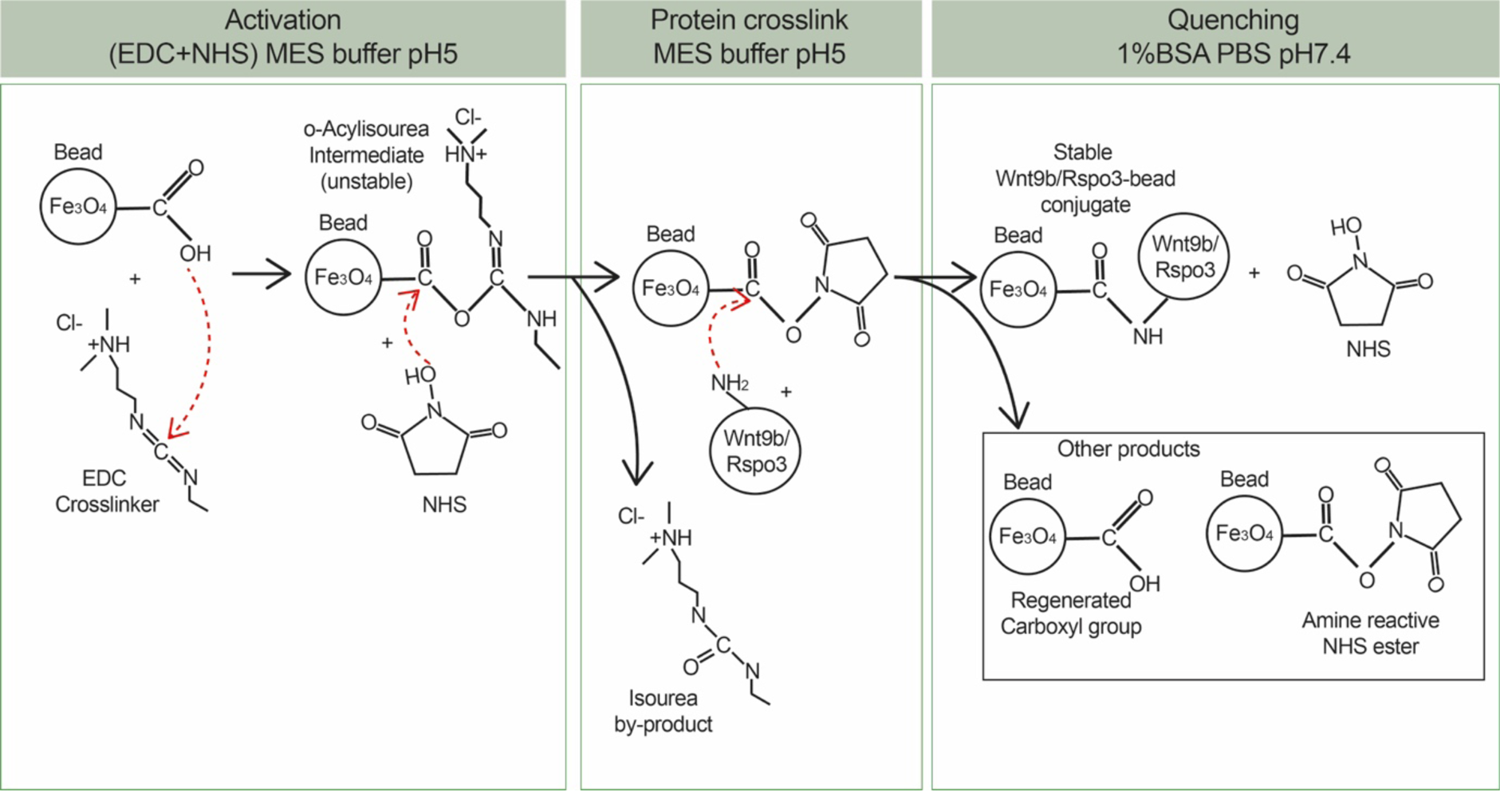
Scheme showing the chemistry behind the conjugation of Wnt9b and Rspo3 proteins onto beads.

**Figure S3.**
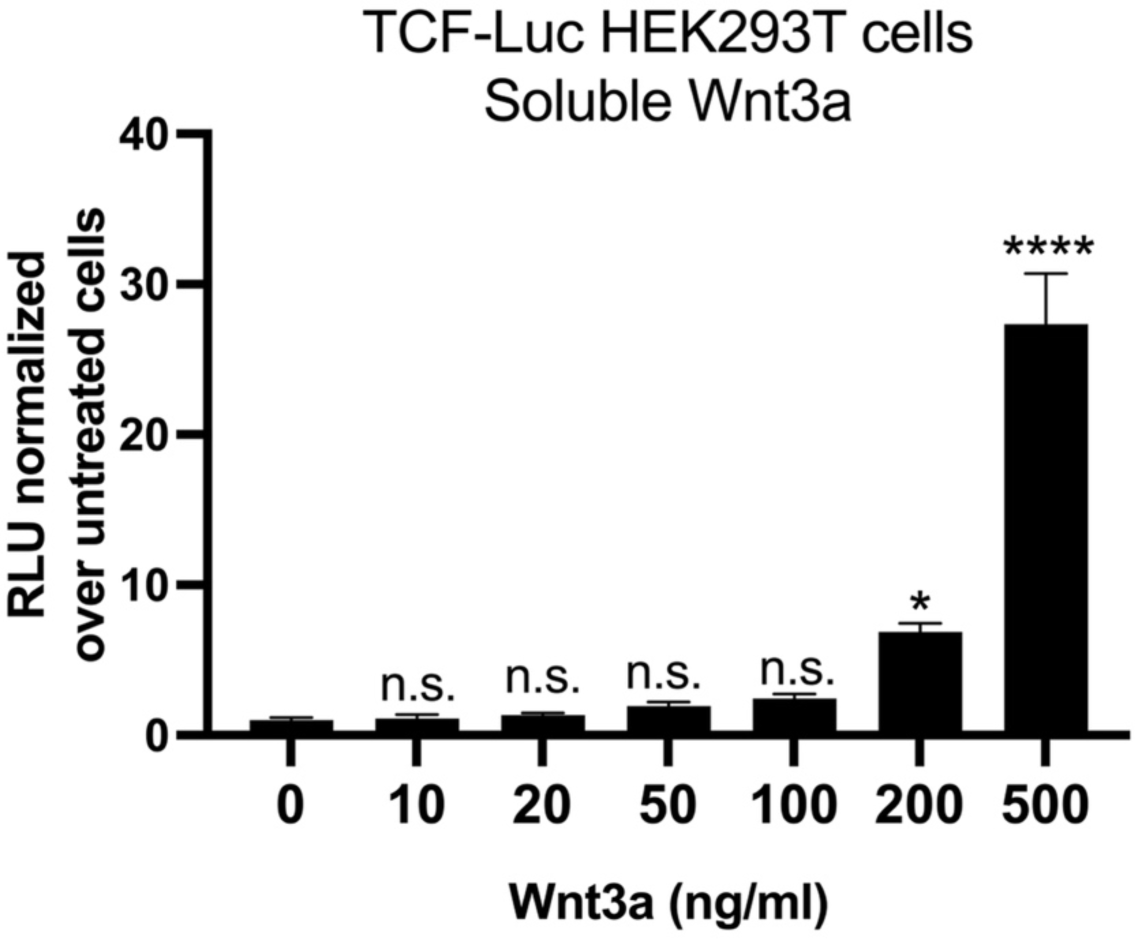
24h response of HEK293TTCF-Luc reporter cells to different concentrations of soluble Wnt3a.

**Figure S4.**
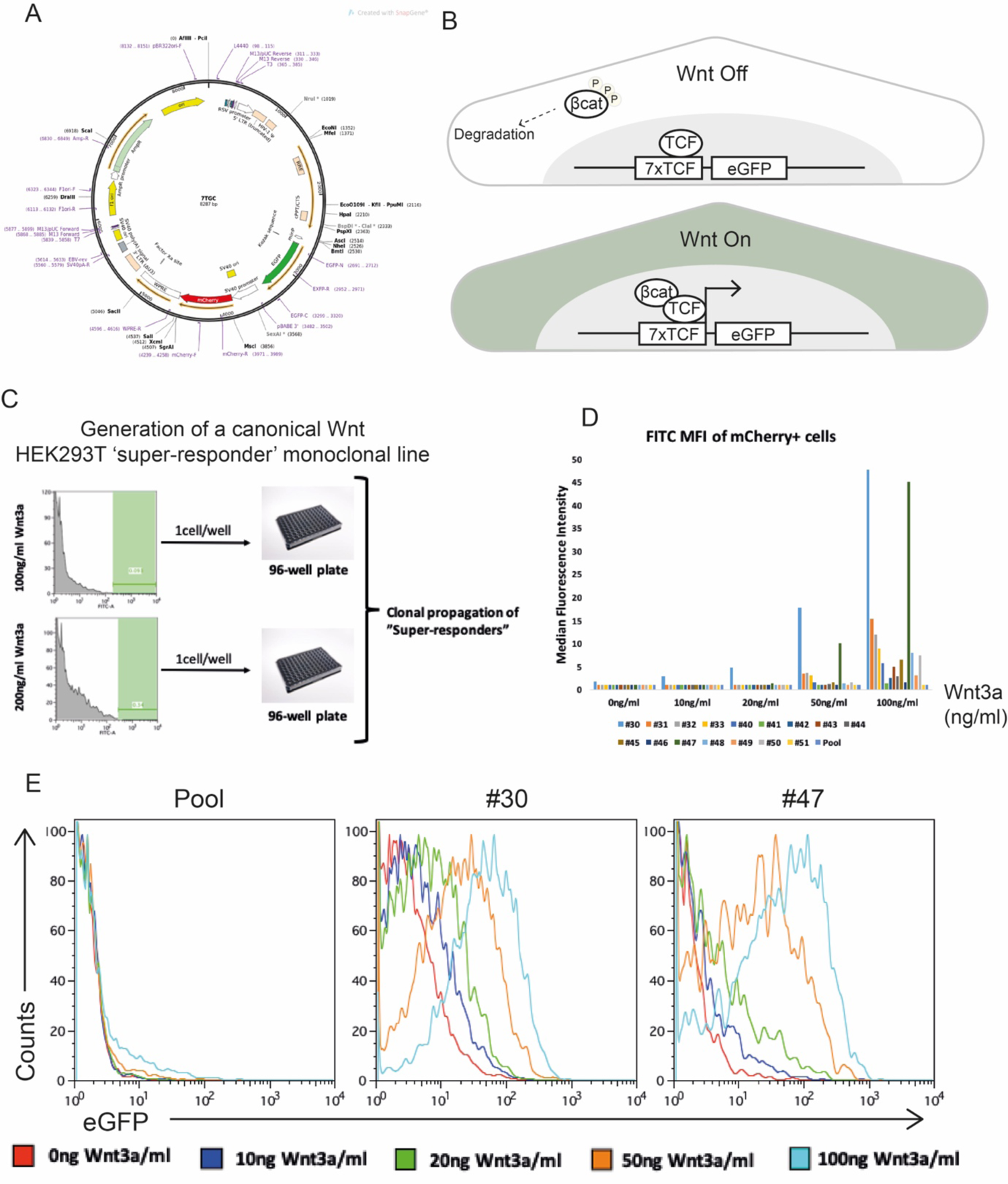
(A) p7TGC plasmid (TCF-eGFP reporter) from Addgene (#24304) contains the sequence coding for eGFP expression under the control of 7x TCF repeats. mCherry is under the control of the SV40 promoter and therefore it’s expression reports successful integration and expression of the plasmid. Image obtained from the provider website. (B) eGFP expression is used as a proxy for canonical Wnt activation levels in ells expressing the TCF-eGFP reporter. (C) Scheme showing experimental strategy for the generation of HEK293T TCF-eGFP canonical Wnt ‘super-responders’. (D) Bar graph shows eGFP median intensity of “Wnt3a Super-responders” upon 24h exposure to 100 ng/ml of Wnt3a. Clones #30 and #47 showed the highest swift in eGFP intensity upon treatment. Most of the clones also showed enhanced responsiveness to purified Wnt3a when compared to the initial pool of successfully transduced cells. (E) Flow cytometry histogram showing responsiveness of the pool of successfully transduced 7pTGC cells, clone #30 and clone #47 to different concentrations of Wnt3a. Both clone #30 and #47 expressed high levels of eGFP upon 100 ng/ml treatment. Clone #47 showed the lowest eGFP basal expression and therefore was selected for further experiments.

**Table S1** Results from RNAseq differential expression analysis performed with Deseq2 (two-by-two comparisons) between W9, W9R3, R3 or CHIR-treated and control PH organoids.

**Table S2** RT-qPCR primer pairs used in this study.

**Table S3** Organoid lines used in this study

## Notes

### Competing Interest Statement

The authors have declared no competing interest.

